# Quantitative live-cell PALM reveals nanoscopic Faa4 redistributions and dynamics on lipid droplets during metabolic transitions of yeast

**DOI:** 10.1101/2020.05.29.123729

**Authors:** Santosh Adhikari, Joe Moscatelli, Elias M. Puchner

**Affiliations:** School of Physics and Astronomy, University of Minnesota, Twin Cities Physics and Nanotechnology (PAN), 115 Union Street SE, Minneapolis, MN 55455, USA

## Abstract

Lipid droplets (LDs) are dynamic lipid storage organelles needed for lipid homeostasis. Cells respond to metabolic changes by regulating the spatial distribution of LDs, as well as enzymes required for LD growth and turnover. Due to LD size below the optical diffraction limit, bulk fluorescence microscopy cannot observe the density and dynamics of specific LD enzymes. Here, we employ quantitative photo-activated localization microscopy (PALM) to study the density of the fatty acid activating protein Faa4 on LDs during log, stationary and lag phases in live yeast cells with single-molecule sensitivity and 30 nm resolution. During the log phase LDs co-localize with the Endoplasmic Reticulum (ER) where the highest Faa4 densities are measured. During transition to the stationary phase LDs translocate to the vacuolar surface and lumen with a ~2-fold increased surface area and a ~2.5-fold increase in Faa4 density, suggesting its role in LD expansion. The increased Faa4 density on LDs is caused by its ~5-fold increased expression level. When lipolysis is induced in stationary-phase cells by diluting them for 2 hrs in fresh medium, Faa4 shuttles to the vacuole through the two observed routes of ER- and lipophagy. The observed vacuolar localization of Faa4 may help activating fatty acids for membrane expansion and reduces Faa4 expression to levels found in the log phase.

## Introduction

Lipid droplets (LDs) are lipid and energy storage organelles of a cell consisting of a neutral lipid core surrounded by a phospholipid monolayer (Tauchi-Sato et al., 2002; Walther and Farese, 2012). Cells maintain their lipid homeostasis and adapt to energy needs by actively regulating the anabolism and catabolism of LDs through regulatory proteins and enzymes (Olzmann and Carvalho, 2019; Walther and Farese, 2012). Fatty acids (FAs) required for the synthesis of neutral lipids such as triacylglycerols (TAGs) as well as FAs released from the breakdown of TAGs by lipases need to be converted into acyl-CoAs by fatty acid activating enzymes (Faergeman et al., 2001). The transition of cells to growth arrest during starvation and back to growth resumption results in major changes in the number, size and subcellular location of LDs. These transitions are mediated by a change in the composition of regulatory proteins and enzymes on LDs. (Kurat et al., 2006; Markgraf et al., 2014; Wang et al., 2014a). Conventional fluorescence microscopy yielded valuable insights into the processes of LD biogenesis, breakdown, and overall co-localization with regulatory enzymes (Kurat et al., 2006; Seo et al., 2017; Wang et al., 2016). However, since conventional fluorescence microscopy cannot resolve LDs and their subcellular location below the optical diffraction limit, little is known about the relation between the exact size of LDs and the density of LD localized regulatory enzymes. Likewise, it is not known how this relation changes upon metabolic shifts. Here we expand our recently developed live-cell single-molecule localization microscopy (SMLM) approach (Adhikari et al., 2019) by resolving LDs at different metabolic states in living yeast cells. For the first time, we gain quantitative insights into the relation between the size of LDs and the density of the endogenously tagged fatty acid activation enzyme Faa4 on LDs upon metabolic transitions. We simultaneously monitor the subcellular localization and redistribution of LDs and Faa4, quantifying their mobility in different metabolic states.

LDs emerge from the ER and maintain a close association with the ER and other organelles for lipid exchange (Jacquier et al., 2011; Schuldiner and Bohnert, 2017). Most of the LD-resident proteins are targeted to LDs during their biogenesis in the ER membrane (Bersuker and Olzmann, 2017; Jacquier et al., 2011). In yeast, the fatty acid activation protein 4 (Faa4) is one of the fatty acyl CoA-synthetases responsible for import and activation of FAs for the synthesis of TAG (Black and DiRusso, 2007; Faergeman et al., 2001). Faa4 as well as the diacylglycerol acyltransferase Dga1, which produces TAGs, are localized to the ER membrane where LDs emerge (Jacquier et al., 2011) and to matured LDs (Kurat et al., 2006; Markgraf et al., 2014).

During nutrient deprivation when yeast cells reach a stationary growth phase, Dga1 and Faa4 dynamically re-localize from the ER to the surface of LDs. In addition Faa4 exhibits increased expression during the stationary phase, which is also crucial for stationary phase survival (Ashrafi et al., 1998; Kurat et al., 2006). The increased amount of Dga1 and Faa4 on LDs causes an increase in local TAG synthesis on LDs and results in their expansion (Markgraf et al., 2014). This expansion of LDs is thought to occur since cells don’t need lipids for membrane expansion during the absence of their growth. In addition, LD-mediated buffering of FAs released during lipolysis reduces lipotoxicity of free FAs (Listenberger et al., 2003). While increased amounts of Faa4 have been observed with conventional fluorescence microscopy on enlarged LDs during the stationary phase, it is unknown if and how cells dynamically control the density of regulatory proteins between the ER and LDs based on metabolic changes.

During the stationary growth phase, some LDs are degraded in the vacuole through micro-autophagy, commonly referred to as lipophagy (Seo et al., 2017; Wang et al., 2014a; van Zutphen et al., 2014). The resulting FAs need to be re-activated by Faa4, which has been shown to enter the vacuole along with LDs during lipophagy (Wang et al., 2014a; van Zutphen et al., 2014). At the same time, LDs at Nuclear Vacuolar Junctions (NVJs) increase their TAG content using the activated FAs (Hariri et al., 2018, 2019). Upon growth resumption from the stationary phase, TAGs stored in LDs are rapidly broken down by LD resident lipases to release and channel DAGs and FAs towards the ER and the vacuole for membrane proliferation (Ganesan et al., 2019; Kurat et al., 2006; Markgraf et al., 2014; Ouahoud et al., 2018). Again, Faa4 is needed to reactivate the released FAs for membrane synthesis.

Previous studies using conventional fluorescence microscopy lack the ability to accurately quantify the spatial distribution of Faa4 and the size of LDs below the optical diffraction limit of ~(250-300) nm. The accurate and simultaneous quantification of LD size, molecular composition, and mobility requires super-resolution microscopy techniques such as PALM (Betzig et al., 2006). In this study, we employ correlative PALM and conventional fluorescence microscopy to study, for the first time, the nanoscopic spatial distribution and the density of endogenously tagged Faa4 on the ER, LDs, and inside the vacuole upon transitions of living yeast cells to different metabolic states. As cells transition to the stationary phase, we find that LDs grow in size with increased Faa4 density on them and get immobilized at the vacuole. Some LDs along with their surface localized Faa4 are eventually taken up by the vacuole where they freely diffuse. When cells in the stationary phase are diluted back to fresh medium for 2 hrs, a few LDs localize again to the ER while Faa4 is predominantly localized to the vacuolar lumen suggesting a mechanism for FA activation inside vacuole and for degrading excess Faa4. Our approach to resolve and quantify the density of regulatory proteins on LDs in living cells may help in future studies to better understand how cells differently regulate their enzyme densities on LDs to maintain lipid and energy homeostasis based on the metabolic needs.

## Results

### Live-cell PALM resolves the sub-cellular localization and dynamics of Faa4 and LDs with ~ 30 nm resolution

Yeast cells actively regulate the spatial distribution, number and size of LDs based on their metabolic state (Wang et al., 2014a; van Zutphen et al., 2014). During the log phase, LDs emerge from and remain closely associated with the ER, where they take up neutral lipids for their expansion. The newly formed LDs are diffraction-limited in size and most of the regulatory proteins are targeted to the surface of LDs during their biogenesis (Kory et al., 2016). Faa4 is one of the fatty acid activating proteins in yeast that has been shown to localize to both the ER and LDs (Kurat et al., 2006; Natter et al., 2005), where its catalytic activity is required to form neutral lipids, phospholipids and protein myristoylation (Ashrafi et al., 1998; Black and DiRusso, 2007). While previous studies using conventional fluorescence microscopy yielded valuable information about the cellular distribution of Faa4, they lack an accurate quantification of the sub-cellular localization of Faa4 and of the size of LDs below the optical diffraction limit. To overcome these limitations, we employed correlative live-cell PALM and conventional fluorescence microscopy. This approach allows us to study the nanoscopic localization and the single-molecule dynamics of Faa4 along with a simultaneous quantification of LD sizes and their motion in living yeast cells. First, we endogenously tagged Faa4 with the photo-switchable fluorescent protein mEos2 in a *S. cerevisiae* strain (W303), which additionally expressed the ER marker Sec63-GFP for correlative conventional fluorescence and PALM imaging. We then recorded a repetitive sequence with one frame of conventional GFP fluorescence at low 488 nm excitation power in the green channel followed by 9 frames with high 561nm laser power and sparse 405 nm photo-activation to detect the single molecule fluorescence of Faa4-mEos2 in the red channel (Supplementary movie1). As expected, LDs in the log phase were localized to the ER but not to the vacuole in co-localization experiments with Sec63-GFP (ER marker) and Ypt7-GFP (vacuole marker) (Fig.1A and 1B). Conventional fluorescence images of Sec63-GFP showed the expected outline of the ER around the nucleus and in proximity to the plasma membrane (Fig 1C, left). The reconstructed PALM images of single Faa4-mEos2 molecules revealed their localization to the ER as well as to LDs (Fig. 1C, middle), which is consistent with previous studies (Kurat et al., 2006; Natter et al., 2005). The localization of Faa4 to LDs was further confirmed by staining LDs with Nile red (Greenspan et al., 1985) (Fig. 1C, right). Importantly, the high ~30nm precision of individual localizations in PALM images estimated with the Thompson formula (Thompson et al., 2002) accurately resolves the size of LDs with a diameter ranging from 100-600 nm (Fig. 1D).

**Fig.1.**
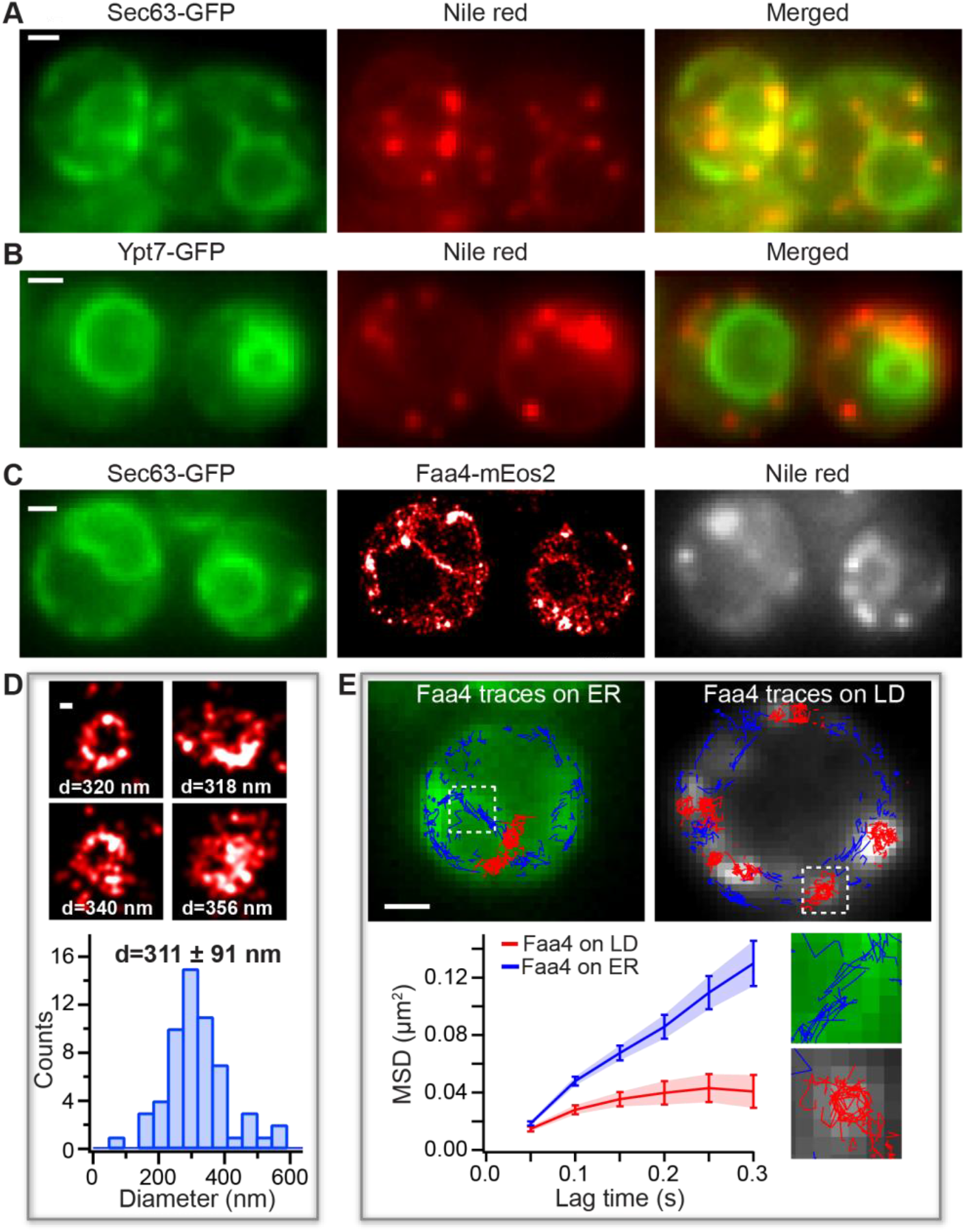
PALM reveals LD sizes and Faa4 dynamics in living cells. (A) The conventional fluorescence images of Sec63-GFP (left), Nile red (middle) and merged (right) show that LDs are co-localized with the ER in the log phase. (B) The conventional fluorescence image of Ypt7-GFP (left), Nile red (middle) and merged (right) indicate that LDs are not localized to the vacuole. (C) Conventional image of Sec63-GFP (left), PALM image of Faa4-mEos2 (middle) and conventional image of LDs stained with Nile red (right) visualize Faa4-mEos2 localizations at the ER and in dense spots that co-localize with LDs. (D) Representative images of super-resolved LDs and histogram of LD diameters with a mean of 311 ± 91 nm (N=58 LDs from 25 cells). Error represents the standard deviation of all LD sizes. (E) Single molecule tracking of Faa4-mEos2 superimposed on Sec63-GFP (left) reveals traces on the ER (blue) and on LDs (red) superimposed on Nile red (right). The MSD of Faa4 traces on the ER (N=3 cells,1270 traces longer than 3 frames) and on LDs (N=8 LDs from 3 cells, 280 traces longer than 3 frames) exhibit free diffusion of Faa4 at shorter lag times but confinement on LDs at longer lag times. Scale bar: 1 μm, zoom: 100 nm.

To gain additional insights into the dynamics of single Faa4 enzymes, we quantified their motion in the ER and on LDs using single-molecule tracking (Manley et al., 2008). The single-molecule tracking of Faa4 revealed its motion along the ER membrane and on the surface of LDs (Fig.1E, upper). The calculated MSD curves showed that Faa4 freely diffuses on ER and on the surface of LDs, but exhibited confinement at longer lag times on LDs (Fig. 1E, lower). The mean diffusion coefficient of all Faa4 enzymes was determined to be D=0.061 ± 0.014 μm^2^/s (Supplementary Fig.1A), which is comparable to other membrane localized proteins (Wu et al., 2015). Since LDs in the log phase are immobile, these results show that Faa4 does not undergo any significant immobilization on LDs compared to the ER.

These results demonstrate that our correlative live cell PALM approach reveals the subcellular localization and the dynamics of Faa4 at the nanoscopic length scale and allows us to quantify the size of LDs in living cells.

### In the stationary phase, Faa4 and enlarged LDs redistribute to the vacuole

When yeast cells exhaust the nutrients in their medium, they enter the stationary growth phase which is characterized by cell cycle arrest along with LD expansion to buffer excess FAs (Hariri et al., 2018; Wang et al., 2014a). During the transition to the stationary phase, LDs have been shown to redistribute to vacuolar micro-domains followed by entry into the vacuole in a process called micro-lipophagy (Seo et al., 2017; Wang et al., 2014a; van Zutphen et al., 2014).

In order to obtain high-resolution insights in the distribution and dynamics of Faa4 and LDs during the transition to the stationary phase, we grew a yeast culture for ~30 hrs without exchange of medium. Co-localization experiments of Nile red with Sec63-GFP and Ypt7-GFP confirmed that LDs redistributed from the ER to either the vacuolar surface or to the vacuolar lumen in these conditions (Fig. 2 A, B). To measure the nanoscopic spatial distribution and the dynamics of Faa4 in this growth phase, we again performed live-cell PALM imaging of Faa4-mEos2. The super-resolved images revealed that Faa4 is localized to bigger LDs in clusters (Fig. 2C, upper). Consistent with previous reports (Wang et al., 2014a) the quantified sizes of LD sizes exhibited an increased diameter (446 ± 81 nm) compared to the log phase (Fig. 2C, lower). In some cells, a fraction of Nile red stained LDs already randomly diffused inside the vacuole (Fig. 2D, upper right, Supplementary movie2). Likewise, the single molecule localizations and traces from Faa4-mEos2 were distributed throughout the vacuolar lumen (Fig.2D, upper left). In order to test if Faa4 was still localized to LDs inside the vacuole, we quantified single molecule tracking of Faa4-mEos2 and single particle tracking of LDs stained with Nile red (Fig.2D, left and right). The MSD of the traces from LDs and Faa4-mEos2 inside the vacuole revealed a similar diffusion coefficient of D=0.10 μm^2^/s (Fig.2D, lower). Since freely diffusing Faa4 would have a diffusion coefficient larger than ~2 orders of magnitude (Swaminathan et al., 1997) and would not be detectable with the used frame rates (Supplementary Fig. 4A, right), this data indicates that Faa4 is still localized to LDs inside the vacuole.

**Fig.2.**
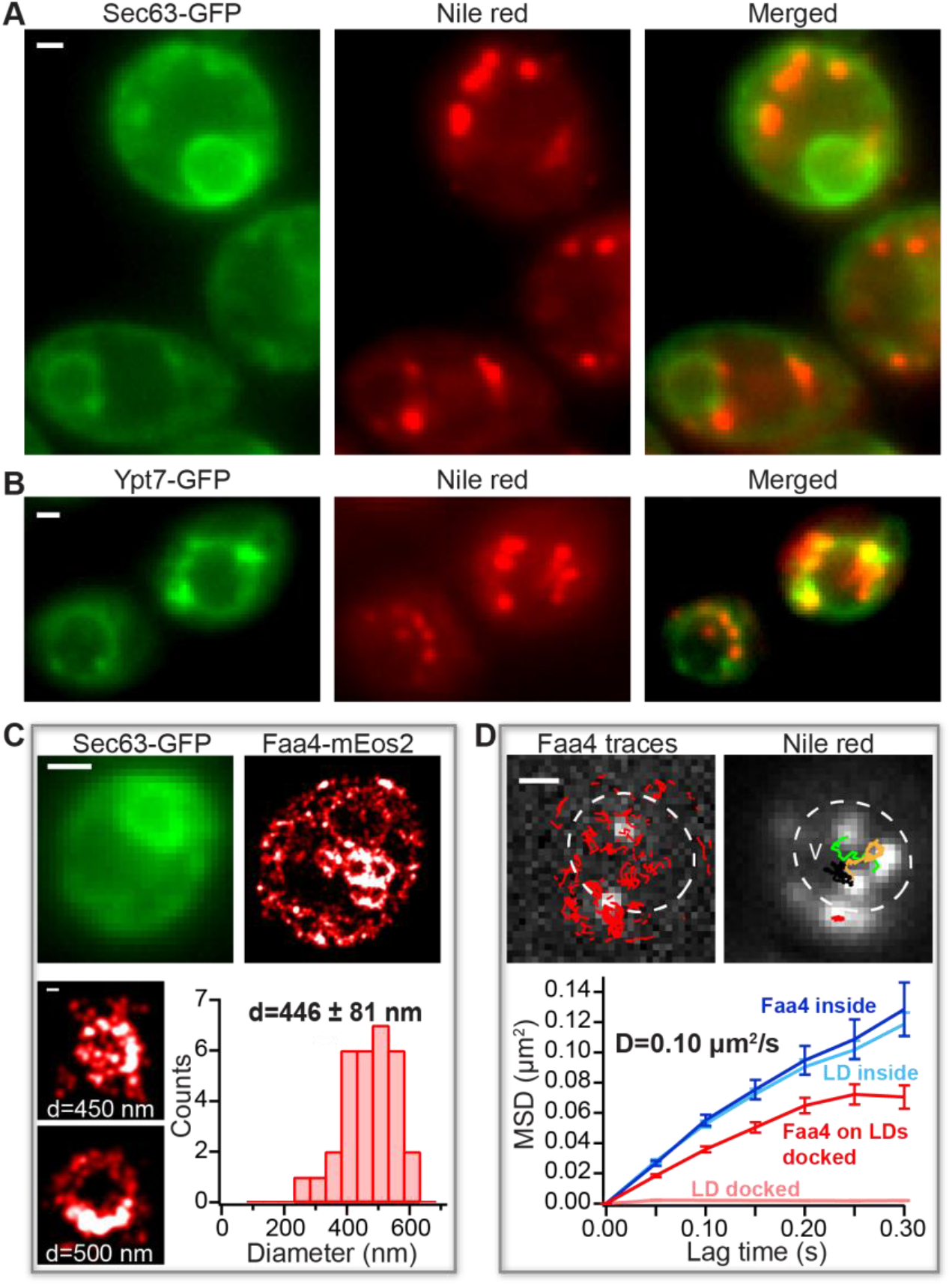
Faa4 redistribution and LD dynamics during the transition to the stationary phase. (A) The conventional fluorescence images of Sec63-GFP (left), Nile red (middle) and merged (right) show LDs not co-localized with the ER during the stationary phase. (B) Conventional fluorescence image of Ypt7-GFP (left), Nile red (middle) and merged (right) reveal LDs either docked to or inside the vacuole. (C) Sec63-GFP (upper, left) visualizes the ER. PALM image of Faa4-mEos2 (upper, right) reveals Faa4 localized to the ER and to LDs. Representative super-resolved images (lower) of LDs with an increased mean diameter of 440 ± 81 nm (N=31 LDs from 10 cells). Error is calculated from the standard deviation of all LD diameters. (D) The single molecule traces (red) superimposed on single molecule signals (white) of Faa4-mEos2 show confined diffusion along the surface of immobile LDs at the vacuole and free diffusion on diffusing LDs inside the vacuole (left). Single particle tracking of LDs with the conventional fluorescence signal from Nile red reveals immobile LDs at the vacuole and free diffusion of LDs inside (right). The MSD of Faa4-mEos2 (N=140 traces from 3 LDs) and LDs (N=425 traces from 3 LDs) exhibit a similar slope with D=0.10 μm^2^/s inside the vacuole (lower). LDs on the vacuole are immobile while the MSD of Faa4 indicates free diffusion with confinement at longer lag times on the surface of LDs as in the log phase (lower). Scale bar: 1 μm, zoom: 100 nm.

Since LDs transition from immobility on the vacuole to random diffusion inside the vacuole (Supplementary movie3), the average size measurement before and the diffusion after the vacuolar entry can be used to calculate the vacuolar viscosity. By using the Stokes-Einstein equation (Saks et al., 2008) with the average diameter of LDs before their entry into the vacuole (440 ± 81 nm, Fig, 2C) as well as the average diffusion coefficient of D = 0.15 ± 0.084μm^2^/s inside the vacuole (Supplementary Fig.2A) the vacuolar viscosity resulted in a value of 6.6 ± 3.9 cP, in agreement with previous studies (Puchkov, 2010). In summary, our correlative conventional fluorescence and PALM approach revealed an increase in LD size upon transition to the stationary growth phase followed by an entry into the vacuole where LDs freely diffuse while Faa4 still localizes to their surface.

### Quantification of live-cell PALM reveals a ~2.5-fold density increase of Faa4 on LDs for their expansion during the stationary phase

During the transition to the stationary phase, yeast cells stop expanding their membranes for cell division and instead increase their TAG content in LDs of increasing size through local synthesis of TAG (Hariri et al., 2018; Kurat et al., 2006). Dga1 catalyzes the final step in the conversion of DAG to TAG with FAs activated by Faa4/Faa1. Consequently, Dga1 re-localizes from the ER to LDs where its activity is needed (Markgraf et al., 2014). During the stationary phase the fatty acid activating enzyme Faa4 has increased expression levels and is also required for stationary phase survival (Ashrafi et al., 1998). When cells are diluted with fresh medium to resume growth (lag phase), TAGs stored in LDs are broken down through lipolysis to release DAG and FAs for the synthesis of other membrane lipids (Kurat et al., 2006; Ouahoud et al., 2018). The catalytic action of Faa4 is required to re-activate FAs released during lipolysis for TAG synthesis on LDs during the stationary phase and for membrane lipid synthesis during the lag phase. During both the stationary and the lag phase, the futile cycle of TAG synthesis and breakdown on LDs has to be broken for a net synthesis or the breakdown of TAGs. One way to break this futile cycle on LDs is a change in density of specific proteins required for TAG synthesis or the breakdown. Hence, we sought to investigate how the density of Faa4 on LDs changes during these metabolic transitions using quantitative PALM.

To establish the conditions of cells in the lag phase, we exchanged the medium of yeast cells in the stationary with fresh medium ~2 hrs prior to imaging. The conventional fluorescence and PALM imaging of LDs and Faa4-mEos2 as well as Sec 63-GFP showed a subcellular distribution of LDs and Faa4 that resembled cells in the log phase. LDs again localized to the ER with a mean diameter of ~312 nm and single Faa4-mEos2 localizations were predominantly found along the ER and on the surface of LDs (Fig.3A). To measure the Faa4 density on LDs, we quantified the surface area and the number of Faa4-mEos2 molecules on individual LDs from the log, stationary and lag phases using their super-resolved images. Counting the number of molecules in protein complexes or organelles with PALM has been a major development in the field and requires methods to correct for blinking artifacts by combining fluorescent bursts emitted from the same fluorophore within a spatio-temporal threshold (Annibale et al., 2011; Deschout et al., 2014; Puchner et al., 2013). However, the application of these methods has so far been limited to fixed cells due to the potential motion of protein complexes or organelles during the data acquisition. Here, we apply molecule counting to live-cell PALM with the following rationales: First, we selected multiple immobile LDs with varying number of mEos2 localizations at low 405nm photo-activation power to avoid spatial overlap of multiple emitters (Supplementary Fig.3A). For each individual LDs, the number of apparent blink corrected molecules at various allowed dark times (t_dark_) was determined by grouping the localizations that appear on the LD within the allowed dark time. This results in dominant double exponential decrease in the number of apparent molecules at shorter dark times due to blinking followed by linear decrease at longer dark times due to false linking (Supplementary Fig.3B). Previous studies with mEos2 have also shown the double exponential distribution of dark times due to the presence of two dark states (Annibale et al., 2010; Lee et al., 2012). By fitting the number of apparent molecules vs. dark time curve with the sum of a double exponential and linear function, we determine the optimal choice for the dark time that balances over counting due to blinking and under counting due to false linking (Supplementary Fig.3B,C). Using this procedure from multiple LDs allows us to determine the average number of localizations from single mEos2 molecules (Supplementary Fig.3D) as in previous studies (Lee et al., 2012) and to convert detected localizations to molecules with an error of 14%. The high degree of correlation (ρ=0.91) between the number of molecules and their localizations from multiple LDs (Supplementary Fig.3D) confirms the accuracy of this approach in quantifying protein count using detected localizations. Second, there may be exchange of bleached and pre-activated mEos2 on LDs with their surrounding. Since our experimental setup employed wide-field excitation and photo-switching, the same fraction of fluorophores is photo-activated and bleached on LDs and their surrounding. Therefore, any bias in over- or underestimating the number of Faa4 molecules due to exchange cancels out.

**Fig.3.**
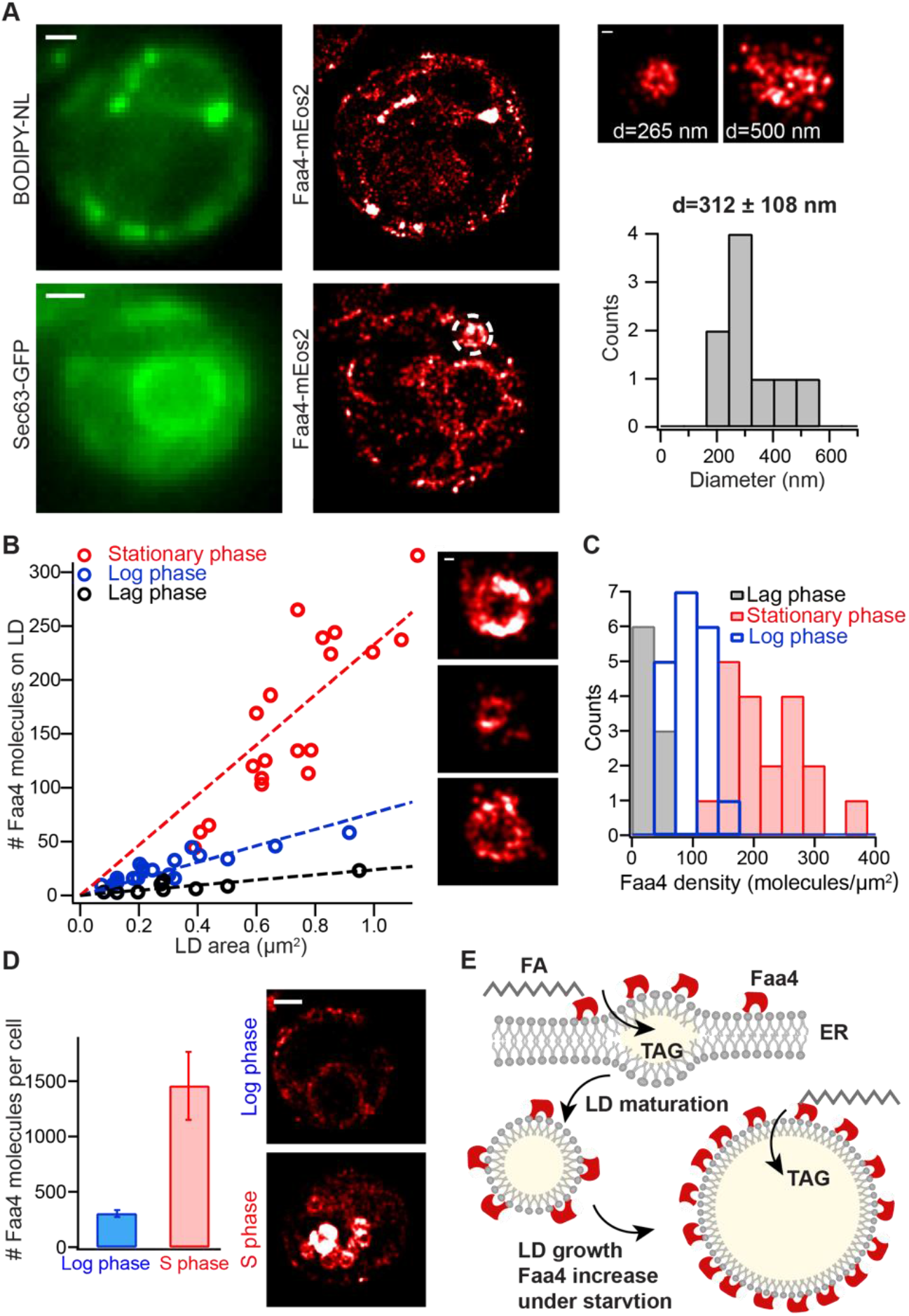
Quantitative live-cell PALM reveals a ~2.5-fold density increase of Faa4 on LDs for their expansion during the stationary phase. (A) In the lag phase single Faa4-mEos2 molecules are localized to LDs visualized with BODIPY-NL (upper) and to the ER visualized with Sec63-GFP (lower). The diameter of LDs with a mean of 312 ± 108 nm (N=9 LDs from 4 cells) is similar to LDs in the log phase (right). (B) The quantified number of Faa4 molecules on individual LDs linearly correlates with their surface area for each growth phase. The larger slope of LDs in the stationary phase compared to log and lag phases indicates an increase in Faa4 density on LDs during stationary phase. The insets show PALM images of LDs in the stationary (upper), log (middle) and lag phase (lower) (C) The histograms of Faa4 densities on LDs shows a roughly 2.5-fold increase of Faa4 density on LDs in the stationary phase compared to log phase. (D) The total number of Faa4 molecules per cell in the stationary phase is roughly 5-fold higher compared to the lag phase and explains the increased Faa4 density on LDs. Error bars are the standard error of the mean calculated using N=9 cells for each case. The representative super-resolution images of Faa4-mEos2 exhibit dense localizations on LDs in the stationary phase and overall much less localizations of Faa4 on the ER and on few emerging LDs in the log phase. (E) A model for LD expansion mediated by a higher density of Faa4 on the LD surface. Scale bars: 1 μm, zooms; 100 nm.

We applied this quantitative PALM approach in each metabolic phase in order to determine the density of Faa4 on individual LDs. The number of Faa4 molecules depended linearly on the surface area of LDs for each growth phase (Fig. 3B). Therefore, the density of Faa4 on LDs in each phase is roughly constant regardless of their surface area. However, the slope of Faa4 molecules vs. LD surface area was different in different growth conditions as seen in the histograms of the respective Faa4 density distributions (Fig. 3C). Log phase cells exhibited a tight distribution with a mean of 95 ± 25 Faa4 molecules/μm^2^, whereas cells in the stationary phase contained on average a ~2.5-fold higher density of Faa4 (219 ± 60 molecules/μm^2^) and a ~6-fold higher number of molecules per LD. The variability of the Faa4 density of LDs in the stationary phase was increased which indicates a more heterogeneous population of LDs compared to the log phase. When cells in the stationary phase were diluted with fresh medium for ~2h and entered the lag phase, the Faa4 density on LDs decreased to values that were lower but similar to cells in the log phase. This observation is consistent with the overall distribution of LDs and Faa4 (Fig. 3A) that resembles the one of log phase cells.

To test if the increased Faa4 density on LDs during the stationary phase is caused by an increased expression of Faa4 or by a regulated increase in its affinity to LDs, we measured the total number of Faa4 molecules per cell under both, the stationary and the log phase. Our results indeed show that stationary phase cells have a roughly five-fold higher number of Faa4 molecules per cell compared to log phase cells (Fig. 3D). Taken together, our quantitative PALM approach reveals a ~5-fold higher expression level of Faa4 during the stationary phase compared to the log phase, which is the predominant mechanism for the measured ~2.5-fold increase in Faa4 density of LDs and its associated increase in catalytic activity for buffering FAs in expanding LDs (Fig.3E).

### Subcellular localization and density of Faa4 during metabolic transitions is dynamically regulated among the ER, LDs and the vacuole for its enzymatic activity

Having gained quantitative insights in the growth phases dependent Faa4 density on LDs, we next quantified the distribution of Faa4 enzymes on the three major organelles where Faa4 is found: the ER, LDs and the vacuole. This measurement of Faa4 redistribution in different growth phases can shed light on where its catalytic activity is need and how the high expression level in the stationary phase is reset to the default values in the log phase.

During the stationary phase, LDs expand at the vacuole through local synthesis of TAGs. Consequently, proteins like Dga1 and Faa4 which are required for TAG and SE synthesis re-localize from the ER to LDs (Kurat et al., 2006; Markgraf et al., 2014) through the cytosol or ER-LD bridges (Kory et al., 2016). The change in relative abundance of proteins on the ER and LDs has therefore been proposed as a regulatory strategy to control their local activity under different metabolic states (Natter et al., 2005). Upon growth resumption from the stationary phase LDs are rapidly consumed through degradation of TAG into DAG and FAs to meet the lipid needs of growing cells (Kurat et al., 2006; Ouahoud et al., 2018). This degradation has been shown to require functional interaction of LDs with the vacuole. The reactivation of released FAs for membrane lipid synthesis is dependent on the enzymatic action of Faa4 (Kiegerl et al., 2019). Therefore, the redistribution of Faa4 among the ER, LDs and the vacuole can be a strategy to direct its catalytic activity to organelles within a cell where it is needed in different metabolic states. To gain insights in the redistribution of Faa4, we quantified the number and the percent of Faa4 molecules per cell that are localized to the ER, LDs and the vacuole in the log-, stationary- and lag phase. Overall, Faa4 was predominately localized to the ER (64%) in the log phase but to LDs (66%) in the stationary phase (Fig 4A, B). This redistribution was accompanied by a dramatic ten-fold increase in the number of Faa4 on LDs compared to the log phase (Fig.4A, B). The number of Faa4 molecules on LDs in the lag phase was roughly two-fold lower compared to the log phase and roughly fifteen-fold lower compared to the stationary phase (Supplementary. Fig.4D). Interestingly, the number of localizations on the ER remained roughly the same in all phases (Fig 4A, B and Supplementary Fig. 4D). In the stationary phase the number and the percent of Faa4 localizations in the vacuole was slightly increased compared to the log phase. However, when stationary phase cells were diluted with fresh medium for ~2 hrs (lag phase), the Faa4-mEos2 signal showed an intense and diffuse fluorescence throughout the vacuole, which was identified by its dark appearance in transmitted light images (Supplementary Fig.4A, B, C). This data indicates a strong accumulation and fast diffusing Faa4-mEos2 inside the vacuole where no LDs were detected any more (Fig 4A, right and Supplementary Fig.4C).

**Fig.4.**
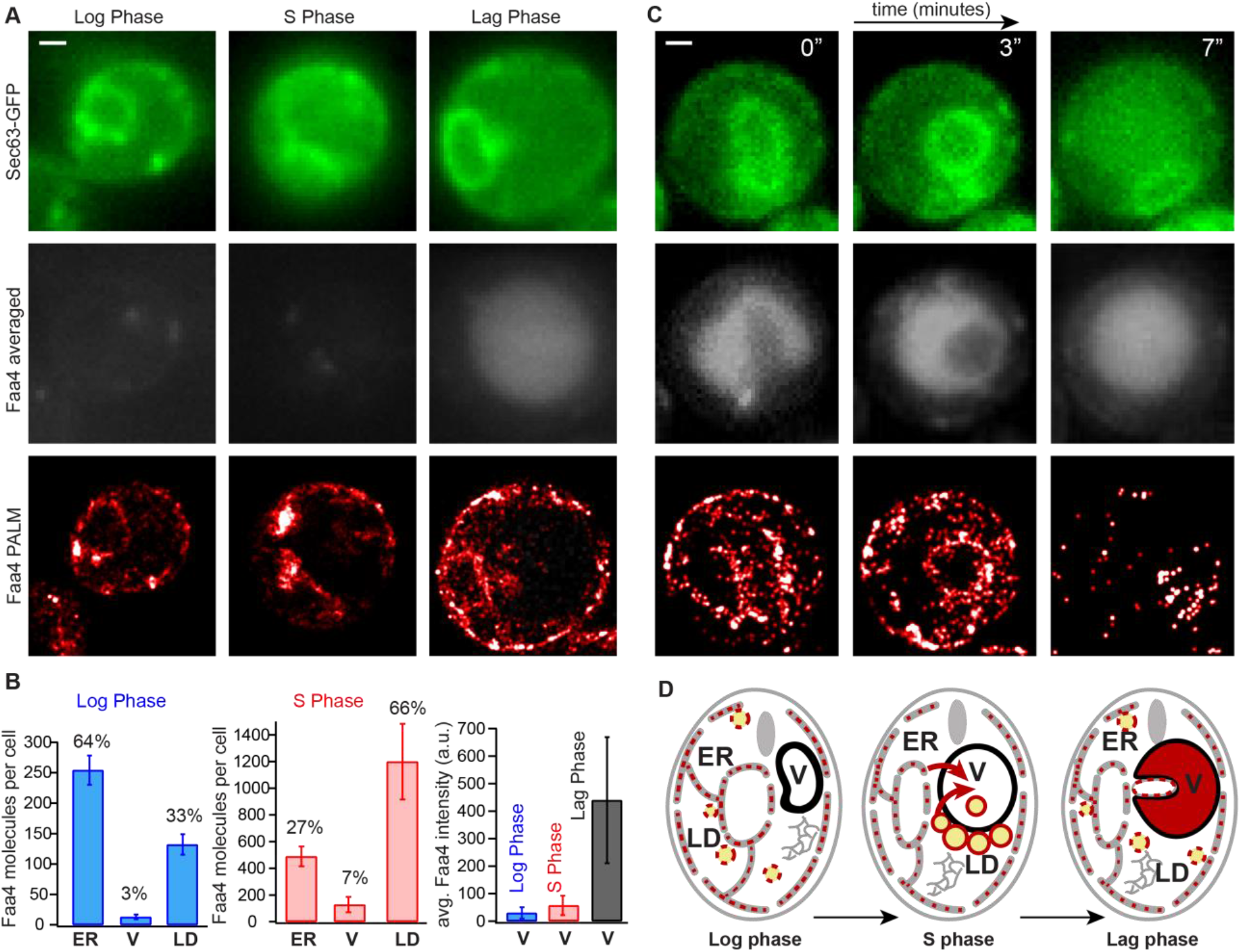
Subcellular Faa4 distribution on the ER, LDs and the vacuole is coupled to the growth phase. (A) Sec63-GFP fluorescence in the log (upper, left), stationary (upper, middle) and lag phase (upper, right) visualizes the ER. The averaged Faa4-mEos2 signal in the log (middle, left), stationary (middle, middle) and lag (middle, right) phases reveals an intense and diffuse Faa4 signal from the circular vacuolar region only in the lag phase. Faa4-mEos2 is similarly localized to the ER and the LDs in the log and lag phase (bottom) but mostly forms dense coats on LDs in the stationary phase. (B) The quantification of the absolute number and the percentage of Faa4 molecules that co-localized with ER, vacuole and LDs in log (N=14 cells) and stationary phase (N=7 cells). Error bars represent standard error on mean. The quantification of the average intensity counts per pixel inside vacuole reveal that Faa4 is predominately localized to vacuole after 2 hrs of dilution in fresh medium (lag phase) (N=5 cells). (C) Time lapse shows how a portion of the Sec63-GFP signal is progressively taken inside the averaged Faa4-mEos2 signal, which indicates that a part ER is taken inside vacuole after 30 minutes of dilution in fresh medium. (D) Schematics showing localization and redistribution of Faa4 among the ER, LDs and the vacuole in the log, stationary and lag phase. The two major routes of lipophagy and ER-phagy shuttle Faa4 into the vacuolar lumen. Scale Bar: 1 μm

The accumulation of Faa4 in the vacuole during the lag phase is likely the mechanism by which cells down-regulate their high Faa4 expression level of the stationary phase to the values observed in the lag phase (Fig. 3D and Supplementary Fig. 4D). So far, our data demonstrates entry of Faa4 into the vacuole at high densities on LDs undergoing lipophagy (Fig. 2D). However, other routes may exist; recently, piece-meal micro-ER-phagy has been shown to maintain proper ER size and protein homeostasis during ER stress recovery in an ESCRT dependent manner (Loi et al., 2019; Schäfer et al., 2019). To test if ER-phagy contributes to the Faa4 re-localization to the vacuole upon growth resumption we imaged the Faa4 distribution ~ 30 minutes after stationary phase cells were diluted in fresh medium. Indeed, we found a significant number of cells that were in the process of or already had taken up parts of the ER into the vacuole, as confirmed by the Sec63-GFP signal (Fig. 4C, Supplementary Fig.4E, Supplementary movie4). Since single molecule localizations of Faa4-mEos2 were detected on the ER inside the vacuole, our data suggests that ER-phagy contributes to Faa4 re-localization into the vacuole upon growth resumption.

## Discussion

Lipid droplets are increasingly recognized as actively regulated organelles that store neutral lipids (TAGs and SEs) in their hydrophobic core (Mejhert et al., 2020; Romanauska and Köhler, 2018). Especially during metabolic transitions of cells, the enzymatic activity of proteins involved in the expansion and breakdown of LDs needs to be actively controlled. Since the density of enzymes is the most fundamental way of tuning the catalytic rate, a quantification of LD size and the number of specific proteins on them is key in understanding the mechanisms of LD metabolism. While previous studies using conventional fluorescence microscopy yield valuable insights into the overall distribution of regulatory proteins on LDs and the dynamics of LDs, they cannot resolve LDs below diffraction limit and count the number of regulatory proteins they contain. To be able to measure protein densities on LDs, quantitative super-resolution microscopy techniques are required that resolve LDs below the optical diffraction limit and enable specific protein counting. In the past most quantitative super-resolution microscopy experiments were performed in fixed cells to avoid movement of the structures under investigation during the long data acquisition time. These experiments employed low 405 nm photo-activation rates to carefully characterize the photo-physical properties of irreversibly bleaching photoactivatable fluorescent proteins such as mEos2. Using this characterization, the fluorescent bursts originating from the same fluorophores were grouped to a photon-weighted position with increased precision and the number of molecules were counted (Durisic et al., 2014; Fricke et al., 2015; Puchner et al., 2013). Here, we transferred this approach to live cell PALM in order to quantify the density of proteins on LDs and to capture their dynamics while cells are transitioning to different metabolic states. Quantitative live-cell PALM is based on two requirements. First, the structures under investigation should not extensively move and should be trackable in the conventional fluorescence channel during data acquisition. This requirement enables grouping of fluorescent bursts from the same fluorophore and the correct assignment of molecules to a structure. Second, the concentration of a protein on a structure must be in equilibrium and not change during the data acquisition time. We note that binding and unbinding of a protein will not affect the outcome of threshold based counting techniques; the same fraction of bound and unbound fluorophores are photo-activated and bleached in a given time. For instance, when 50% of fluorophores on a LD have been imaged and bleached, 50% of unbound fluorophores will have been bleached as well. Therefore, binding and unbinding will not alter the fraction of unbleached fluorophores and the quantification of their number. The linear relation (ρ=0.91) between the number of localizations and the number of blink-corrected molecules on individual LDs demonstrates that the number of detected localizations is an accurate measure for quantifying protein numbers as previously demonstrated in other systems (Lee et al., 2012) and confirmed with intracellular calibration standards (Puchner et al., 2013). In this study, only LDs that freely diffused inside the vacuole during starvation violated the first requirement of immobility. All other LDs were immobile to determine their size and the number of associated proteins. Since the transition of cells to different metabolic states and the associated change in density of regulatory proteins takes place on the timescale of hours (Brauer et al., 2008), potential density changes during the data acquisition time of several minutes can also be considered insignificant.

In this study we applied quantitative live cell PALM to Faa4, one of the fatty-acyl CoA synthetases required to activate FAs for the expansion of LDs (synthesis of neutral lipids and other complex lipids) as well as for their breakdown (beta-oxidation). Previous studies using conventional bulk fluorescence of Faa4-GFP followed LD dynamics and the overall distribution of Faa4 to show that Faa4 dynamically localizes among the ER, LDs and the vacuole based on the nutritional state of a yeast cell (Kurat et al., 2006; Natter et al., 2005; Wang et al., 2014a; van Zutphen et al., 2014). However, these studies lacked an accurate quantification of Faa4 abundance, LD size, and Faa4 densities on LDs, a fundamental quantity in determining catalytic activity.

In the log phase, we observed exclusively immobile LDs co-localized with the ER, demonstrating the LD biogenesis and expansion consistent with previous studies. The super-resolved images of LDs with Faa4-mEos2 further revealed the size distribution of LDs with a mean diameter of 310 nm, which agrees with previous EM size measurements (Wang et al., 2014b). Most importantly, our quantitative live cell PALM approach enabled us to quantify the number of Faa4 enzymes and their diffusion on the ER, LDs, and the vacuole. In the log phase, more than 60% of Faa4 localizes to the ER membrane and about 34% to LDs, which are the subcellular locations where the catalytic activity of Faa4 is needed for the activation of FAs and their subsequent synthesis into neutral lipids. The average diffusion coefficient of Faa4 on the ER of 0.061 ± 0.014 μm^2^/s agrees with the published diffusion coefficient of FAs measured using BODIPY-C12 (Adhikari et al., 2019) and indicates that Faa4 freely diffuses in the ER membrane. Similarly, we found that Faa4 on LDs freely diffuses on the surface of LDs as also observed with the LD localized protein Dga1 using FLIP (Jacquier et al., 2011). These results suggest that Faa4 is homogeneously distributed in the ER membrane and targeted to emerging LDs through passive diffusion from the ER membrane.

During starvation and the transition to the stationary phase, LDs have been shown to expand by accumulating neutral lipids followed by lipophagic degradation in the vacuole (Markgraf et al., 2014; Wang et al., 2014a; van Zutphen et al., 2014). Consistent with these findings, our data showed LDs either docked to the vacuole or already diffusing inside the vacuole. Notably, our quantified average diameter of LDs (440 nm) in the stationary phase was significantly larger compared to the 310 nm in the log phase and corresponds to a ~2-fold increase in surface area and ~3-fold expansion of their volume. The unique capability of PALM to accurately measure the size and to estimate the number of Faa4 molecules on LDs further revealed a ~2.5-fold increase in Faa4 density on LDs compared to the log phase. This density increase of Faa4 might therefore serve to activate FAs for LD expansion in the stationary phase. In fact, Dga1, responsible for converting DAG to TAG by adding one fatty acid activated by Faa4, has been shown to exclusively localize to LDs during the stationary phase (Markgraf et al., 2014). The linear correlation of the Faa4 numbers with the surface area of LDs and the overall increase in Faa4 expression level suggests that the Faa4 density is passively regulated through its abundance rather than actively recruited through changes in enzyme affinity. This suggestion agrees with a previous study about the role of an increased Faa4 expression level for cell survival during the stationary phase (Ashrafi et al., 1998).

Our tracking data revealed that LDs docked to the vacuole are immobile while single Faa4 molecules still freely diffuse on the surface of LDs. However, LDs exhibit random diffusion inside the vacuole with an average diffusion co-efficient of D=0.15 μm^2^/s. The similar diffusion coefficient of Faa4 indicates that Faa4 is still localized to LDs that entered the vacuole through lipophagy (Tsuji et al., 2017; Wang et al., 2014a). The viscosity inside the vacuole estimated using Stokes Einstein with the determined diffusion coefficient of LDs (D=0.15 μm^2^/s) and their average diameter (440 nm) was 6.6 ± 3.9 cP, which is in agreement with previous measurements (Puchkov, 2010, 2012).

When cells in the stationary phase are diluted back to fresh medium, LDs are rapidly consumed for membrane expansion during cell proliferation. Previous studies have shown that LD resident lipases (Tgl3) are required for the lipolysis of TAGs to generate DAG and FAs (Ganesan et al., 2019; Kurat et al., 2006; Ouahoud et al., 2018). The resulting FAs in turn need to be activated by fatty acid activating enzymes to serve together with DAG as building blocks for the synthesis of membrane phospholipids. In our study, we observed almost no LDs around or inside the vacuole, indicating their rapid degradation upon growth resumption. Therefore, our detected dramatic increase of Faa4 inside the vacuole may serve the activation of FAs released from LD breakdown in the vacuole. Since cells in the log phase have a five-fold lower expression level of Faa4, the localization of Faa4 to the vacuole could therefore in addition contribute to downregulating Faa4 expression levels after growth resumption (Fig.4E). Our data supports two mechanisms how Faa4 enters the vacuole. The first mechanism is the dramatic 10-fold increase of Faa4 on LDs in the stationary phase and its subsequent uptake into the vacuole during lipophagy. The second mechanism is ER-phagy, which may account for the two-fold decrease of ER-localized Faa4 in the log phase compared to the stationary phase.

In summary, we presented the first study that employs PALM in living cells to quantify the size of LDs and their associated Faa4 numbers and mobility throughout different metabolic states. Our results demonstrate that LD expansion and breakdown during metabolic transitions is coupled with density changes of Faa4 on LDs. This is consistent with the required catalytic activity of Faa4 on LDs in each metabolic state. The 2.5-fold increase in Faa4 density on LDs in the stationary phase is functionally correlated to their increased size and caused by increased expression level. In future studies our quantitative live-cell PALM approach can be transferred to numerous other regulatory proteins. The resulting high-resolution data may give new insights into the regulation of LD biogenesis, growth and mobilization in a cell’s normal physiological condition and how such regulation might be altered in metabolic disorders.

## Materials and Methods

### Yeast strains

The endogenous tagging of Faa4 with mEos2 in a W303 *S. cerevisiae* strain as well as the chromosomal integration of Sec63-GFP was described previously in reference (Adhikari et al., 2019). The W303 yeast strain with integrated Ypt7-GFP was described in reference (Puchner et al., 2013).

### Sample Preparation

Yeast strains were grown overnight in synthetic complete dextrose (SCD) medium with shaking (270 rpm) at 30 °C. The next morning, cells were diluted 1:50 in SCD to an OD of ~0.15 and allowed to grow for 4 hrs to the log phase with and OD of 0.5. For cells in the late diauxic-shift/stationary phase, cells were grown for ~30 hrs without exchange of medium. To prepare cells in the lag phase, stationary phase cells (~30 hrs in same medium) were diluted with fresh SCD to an OD of ~0.2 and imaged after ~2 hrs. Cells in Fig.4C and supplementary Fig.4E were imaged after diluting stationary phase cells in fresh media for ~30 minutes to observe Faa4 distribution at shorter lag time. To adhere cells for imaging, the chambered borosilicate coverglasses (eight well, Lab-Tek II; Sigma-Aldrich) were incubated with ~75 μL sterile filtered Concanavalin A (ConA, Sigma-Aldrich) at a concentration of 0.8 mg/ml in diH2O for 30 mins. After washing the coverglasses three times with deionized water (Millipore), 300 μL of yeast cells were incubated on the coverglass at an optical density (OD) of ~0.12 and allowed to settle for ~30 mins. For lipid droplets (LD) staining with Nile red (Thermo Fisher) or BODIPY-NL (Thermo Fisher), dyes were directly added at a concentration of 100 nM and imaged after ~10 mins.

### Experimental setup and data acquisition

All conventional fluorescence and PALM data were acquired on a Nikon Ti-E inverted microscope with a Perfect Focus System. Four excitation lasers (405, 488 and 561 nm, OBIS-CW; Coherent) were combined using dichroic mirrors. The laser beams were aligned, expanded and focused to the back focal plane of the objective (Nikon-CFI Apo 100x Oil immersion N.A 1.49). The power and shutters of lasers were computer controlled using the Hal4000 software (Zhuang lab, Harvard). A quad-band dichroic mirror (zt405/488/561/640rdc; Chroma) was used to separate the fluorescence emission from the excitation light. For two-color imaging the red (595 nm) and green (525 nm) fluorescence were split by a dichroic longpass beam-splitter (T562lpxr BS; Chroma) followed by bandpass filters ET525/50 (Chroma) in the green channel and ET595/50 (Chroma) in the red channel respectively. The fluorescence emission was recorded at the frame rate of 20 Hz on an electron multiplying CCD camera (Ixon89Ultra DU-897; Andor) cooled to −68 °C and with a gain and pre-amp gain of 30 and 5.1 respectively.

The 405 nm laser was used to photo-activate mEos2 molecules at a power density of 0.1-1 Wcm^−2^ whereas the 488 nm laser was used to excite GFP, BODIPY-NL (493/503) and Nile red molecules at a power density between 0.07 and 0.7 Wcm^−2^ for conventional fluorescence imaging. The excitation of single photo-activated mEos2 molecules for PALM imaging and single molecule tracking was performed with a 561 nm laser at a power density of 0.8-1 kWcm^−2^. Since Nile red can also be excited at 561 nm, we added Nile red after imaging mEos2 in the Sec63-GFP+Faa4-mEos2 yeast strain. We note that pre-activated mEos2 has been shown to have a 10-fold lower fluorescence intensity compared to GFP (Puchner et al., 2013) and accordingly we did not observe any significant leaking into the GFP channel. For the simultaneous imaging of GFP and Nile red, both fluorophores were excited with the 488 nm laser and the fluorescence of GFP was detected in the 525 nm emission channel and Nile red in the 595 nm channel. BODIPY-NL was added and imaged after Sec63-GFP imaging or imaged at a ~10 fold lower 488 nm excitation intensity compared to GFP imaging to avoid potential crosstalk with the GFP signal.

### Camera calibration

The EMCCD camera was calibrated to obtain to gain (e/ADU) and the camera offset. For the calibration, camera pixels were exposed to varying intensity of light with an EMCCD gain of 30. The slope of the mean intensity vs variance graph accounting for the extra noise factor (F^2^) was used to calculate the gain. The calculated gain of 0.166 e/ADU and the offset of 200 counts matched the manufacturers’ specification at EMCCD gain 30 and pre-amp gain 5.1.

### Conventional fluorescence image analysis

The conventional fluorescence images of Sec63-GFP, Ypt7-GFP, BODIPY-NL and Nile red were generated by averaging (50-100) image frames. The conventional fluorescence images of Faa4-mEos2 were obtained by averaging (20-60) 561 nm excitation frames.

### Super-resolution and single-molecule tracking data analysis

PALM data analysis was performed using the INSIGHT software (Zhuang lab, Harvard) that uses a Gaussian PSF model (Gaussian heights ≥ 50 photons, width (260-650) nm, ROI: 7×7 pixels with pixel size 160 nm) to fit the single molecule signal. The obtained molecule list consisting of single molecule co-ordinates, frames of appearance, widths, heights and total counts were exported for the further analysis described below. Super-resolution images were generated by rendering single molecule localizations as 2D Gaussians whose width is weighted by the inverse square root of the detected number of photons.

For single molecule tracking analysis, the single molecule localizations that appeared within 0.48 μm in consecutive frames (50 ms exposure) were linked to a trace. The average distance between activated mEos2 molecules in a frame was kept low by adjusting the photo-activation density so that different molecules are not accidentally linked. Only single-molecules traces lasting for at least 3 frames (20 Hz) were extracted for further analysis. From each trace, the Mean Squared Displacement (MSD) for a particular lag time Δt was calculated by averaging the squared displacements over all time intervals of length Δt with a custom written Igor-pro program. The mean diffusion coefficient (D) was then calculated by linear fitting the averaged MSD vs Δt curve from more than 100 traces with MSD= 4D Δt +2σ^2^ and localization precision σ.

### Single particle tracking (SPT) of LDs

The fluorescence signals from the neutral lipid core of LDs stained with Nile red or BODIPY-NL were detected with INSIGHT as diffraction limited blobs and fitted with Gaussians to determine the centroids. The obtained localizations were linked to traces and further quantified as described in the previous section about single-molecule tracking.

### Blink correction and estimate of Faa4 molecules

In order to convert the number of localizations to the number of molecules, we employed a spatio-temporal threshold method for grouping localizations from the same mEos2 molecule that has been applied and confirmed in fixed cells (Durisic et al., 2014; Puchner et al., 2013). These methods are based on the fact that mEos2, once photoactivated, blinks for a variable time with various dark-times between fluorescent bursts until it irreversibly bleaches (Supplemental Figure 3A). The low 405 nm photoactivation power used throughout PALM experiments results in significantly longer times between the photoactivation of nearby mEos2 molecules compared to the time between fluorescent bursts from the same fluorophore. Therefore, fluorescent bursts appearing within the spatial resolution and a maximum dark-time can be grouped to determine the blink-corrected number of molecules.

To determine the optimal dark-time in this study, localizations from individual LDs at low 405 nm photo-activation power were selected for further analysis (Supplementary Fig.3A, right). For each individual LD, the number of apparent blink corrected molecules at various allowed dark times td was determined by grouping localizations appearing within 650 nm and the allowed td (Supplemental Figure 3B, left). The spatial threshold of 650 nm is the maximum diameter of LDs and accounts for diffusion of photoactivated mEos2 along the surface of LDs. At td=0 s localizations that appear in consecutive frames are grouped and the number of apparent molecules therefore corresponds to the number of fluorescent bursts. As td increases, more and more fluorescent bursts from the same mEos2 molecules are grouped which results in a double exponentially decay of the apparent number of molecules and dominates at shorter dark times. This double exponential decay is caused by two previously identified dark states with different decay times (De Zitter et al., 2019; Lee et al., 2012). At longer dark times when most fluorescent bursts are incorrectly grouped, the probability for falsely grouping two mEos2 molecules keeps increasing and results in a dominating linear decrease in the number of apparent molecules. Therefore, the obtained apparent molecules vs td curve for each individual LD was fitted with the sum of a double exponential and linear function (Supplemental Figure 3B, left). For each LD, the optimal td was determined to be where the fraction of over counting balances the fraction of under counting due to false grouping (Supplementary Fig.3 B, C). The average number of molecules per localizations of 0.13 ± 0.04 from N=9 different LDs (Supplementary Fig.3D) was then used for all PALM data to convert localizations to molecules and to quantify the Faa4 density on LDs (Fig.3B,C), Faa4 molecules per cell (Fig.3D) and Faa4 molecules on the ER, LDs and in the vacuole (Fig.4B) with an error of 14%. While the number of Faa4 molecules on immobile LDs can directly be determined by blink correction, the free diffusion of Faa4-mEos2 on the ER and in the vacuole prevents reliable blink correction and makes a localization-based approach more comparable and reliable in different cellular compartments. We note that previous reports in fixed cells found that only ~60% of mEos2 molecules are detected in PALM experiments (Durisic et al., 2014; Puchner et al., 2013). While this observable fraction has not been determined so far in living cells due to the experimental challenges, the true number of molecules is likely larger than the number of detected molecules. However, both uncertainties for converting localizations to the number of molecules only affect the absolute numbers and do not bias the relative densities or abundances that are relevant for this study.

### 405 nm photo-activation energy correction

We found that the expression level of Faa4-mEos2 is ~5-fold higher in stationary phase compared to log phase (Fig.3D). In order to maintain a constant and non-overlapping detection of mEos2 molecules in each data acquisition frame, this difference in expression levels makes it necessary to acquire PALM data with different 405 nm photo-activation energy densities. As a result, different fractions of total mEos2 molecules can be activated in different movies based on the delivered 405 nm energy. Therefore, it is necessary to verify that the fraction of activated molecules only depends on the delivered 405 nm energy but is independent of acquisition time and mEos2 expression level. Once a calibration curve relating the fraction of activated mEos2 molecules and the delivered 405 nm is obtained, the difference in the fraction of activated mEos2 can be adjusted to compare the localizations across different movies using the delivered 405 nm energy.

In order to verify that the fraction of activated mEos2 molecules only depends on the delivered 405 nm energy, we analyzed PALM data from cells in the log and stationary phase that were imaged with different 405 nm powers and data acquisition times. From the obtained mEos2 localizations for both the log and stationary phase, the fraction of activated mEos2 molecules was calculated as a function of data acquisition time. As expected, the fraction of activated mEos2 molecules is lower for stationary phase cells at a particular data acquisition time since a longer total data acquisition time is required for 100% activation (Supplementary Fig.3F). Next, we calculated the cumulative mEos2 localizations vs delivered 405 nm energy using the 405 nm power and exposure time (Supplementary Fig.3G). The resulting cumulative fraction of activated mEos2 molecules vs 405 nm energy overlapped for both log and stationary phase cells and verifies that the fraction of detected mEos2 molecules only depends on the delivered 405 nm energy (Supplementary Fig.3H).

To quantify Faa4 densities on LDs in Fig.3 B, C, recorded at different 405 nm energies, the delivered 405 nm energy was calculated by multiplying the 405 nm power with the exposure time of each frame and integrating over time. For LDs from movies with a total delivered energy less than ~5 mJ (95% photo-activation), the number of localizations were adjusted to 100% activation using the percent activation vs energy calibration curve (Supplementary Fig.3H). For the quantification of Faa4 molecules in Fig.3D and Fig.4B, the energy delivered in each movie corresponded to more than ~95% activation. Therefore, no adjustments for the number of localizations were used.

### Radius and Faa4 density of LDs

In order to determine the radius of LDs, molecule lists of Faa4-mEos2 belonging to individual LDs were first extracted. The radius r of LDs was then approximated by computing the average standard deviation in x and y assuming LDs are spherical in 3D. The surface density of Faa4 on LDs was then obtained by calculating the ratio of the number of Faa4 molecules and the surface area of a sphere of radius r.

### Subcellular localizations of Faa4

To quantify the subcellular localizations of Faa4 on the ER, LDs and the vacuole in different metabolic states, Faa4-mEos2 localizations from super-resolution images were first transformed to and superimposed on the conventional (GFP) channel in the INSIGHT software. This third order polynomial transformation was determined and verified by localizing and mapping TetraSpeck microspheres (Invitrogen T7279) which fluoresce in both channels. The number of Faa4 localizations that co-localized with the bright Nile red puncta (LD marker) and Sec63-GFP (ER marker) were determined from superimposed super-resolution and conventional images in INSIGHT. In the super-imposed images, the dense circular Faa4-mEos2 structures always co-localized with the Nile red signal. To further increase the statistics, dense circular structures in super-resolution images of Faa4-mEos2 in cells with no Nile red/BODIPY-NL staining were therefore also considered to be from LDs and included. The remaining localizations in each cell that did not co-localize with the ER marker Sec63-GFP were considered to be from the vacuole. While vacuolar Faa4-mEos2 was detected as single molecule localizations in the log and stationary phase due to its slow diffusion, the fast vacuolar diffusion of Faa4-mEos2 in the lag phase prevented the detection of single molecules. However, the signal of the freely diffusing Faa4-mEos2 in the vacuolar lumen was evident from a homogenous and diffuse florescence. To compare the relative abundance of Faa4 in the vacuole in different growth conditions (Fig. 4A) we averaged ~20 frames of the 561 nm channel and used the CCD counts per pixel as a metric for its relative abundance.

## Acknowledgements

We thank Doug Mashek and his lab as well as Christer Ejsing and Florian Froehlich for helpful discussions. From the Puchner lab, we thank Maria-Paz Ramirez Lopez for making the Sec63 yeast strain. Research reported in this publication was supported by the National Institute of General Medical Sciences of the National Institutes of Health under Award number R21GM127965

## Supplementary Material

**Suppl. Fig.1:**
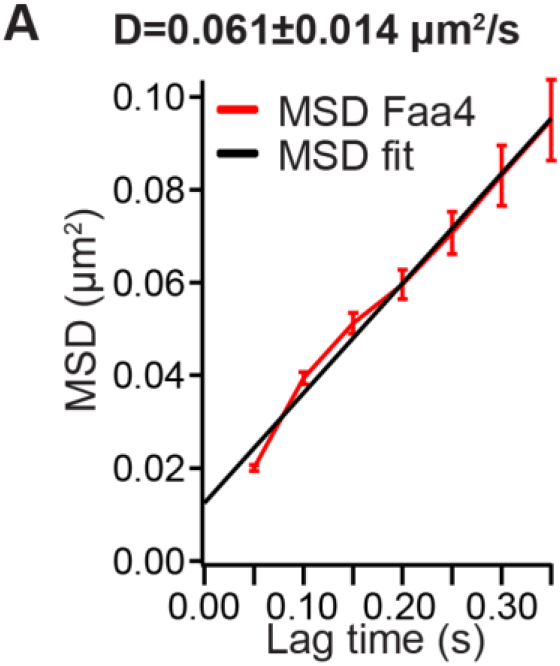
Single-molecule tracking quantifies Faa4 mobility. (A) Fitting the average MSD plot of Faa4 traces on the ER and on LDs with a function results in a diffusion coefficient of D=0.061± 0.014 μm^2^/s. The averaged MSD is calculated from Faa4 traces in 5 cells each containing more than 100 traces lasting at least 3 acquisition frames recorded at 20 Hz. The error of the diffusion coefficient is calculated using the standard deviation of diffusion coefficients from 8 cells.

**Suppl. Fig.2:**
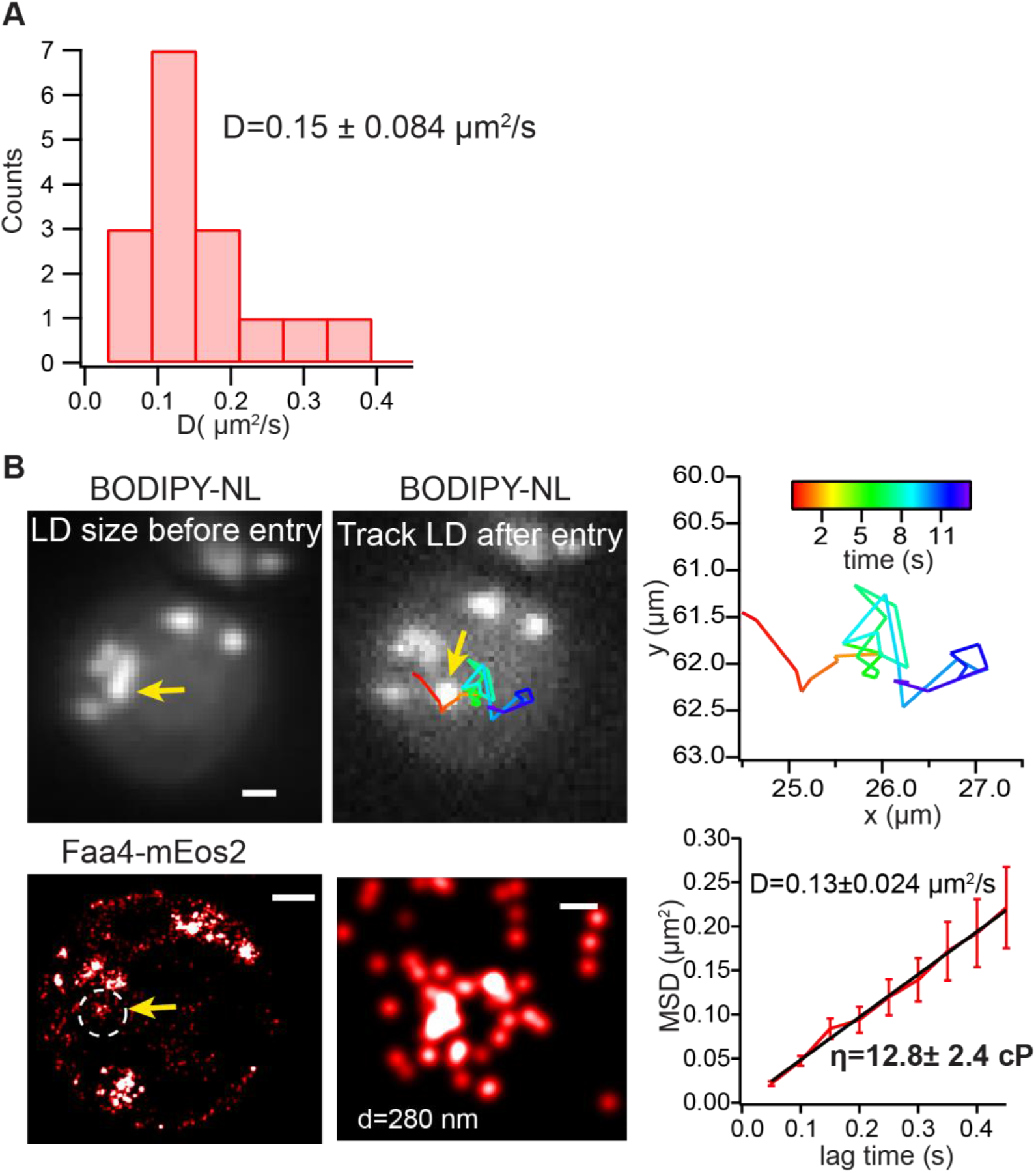
Single particle tracking measures LD diffusion inside the vacuole in the stationary phase. (A) Histogram of LD diffusion coefficients calculated from N=16 LDs and 5 cells with a mean of 0.15 ± 0.084 μm^2^/s. The error of the diffusion coefficient is calculated using the standard deviation of diffusion coefficients from all LDs. (B) Conventional fluorescence image of BODIPY-NL (top, left) shows LDs in cluster. (Bottom, left) PALM image of LDs clustered around the vacuole. Zoom shows a super-resolved LD with a radius of d=280 nm before entry into the vacuole. (Top, right) After the LD marked by an arrow enters the vacuole, it randomly diffuses in the vacuolar lumen with a diffusion coefficient of D=0.13 ± 0.024 μm^2^/s (bottom, right). The vacuolar viscosity of 13 ± 2.4 cP was calculated with an LD size of d=280 nm and diffusion coefficient of D= 0.13 ± 0.024 μm^2^/s at room temperature using the Stokes-Einstein equation. Scale bar: 1 μm, zoom: 100 nm.

**Suppl.Fig.3:**
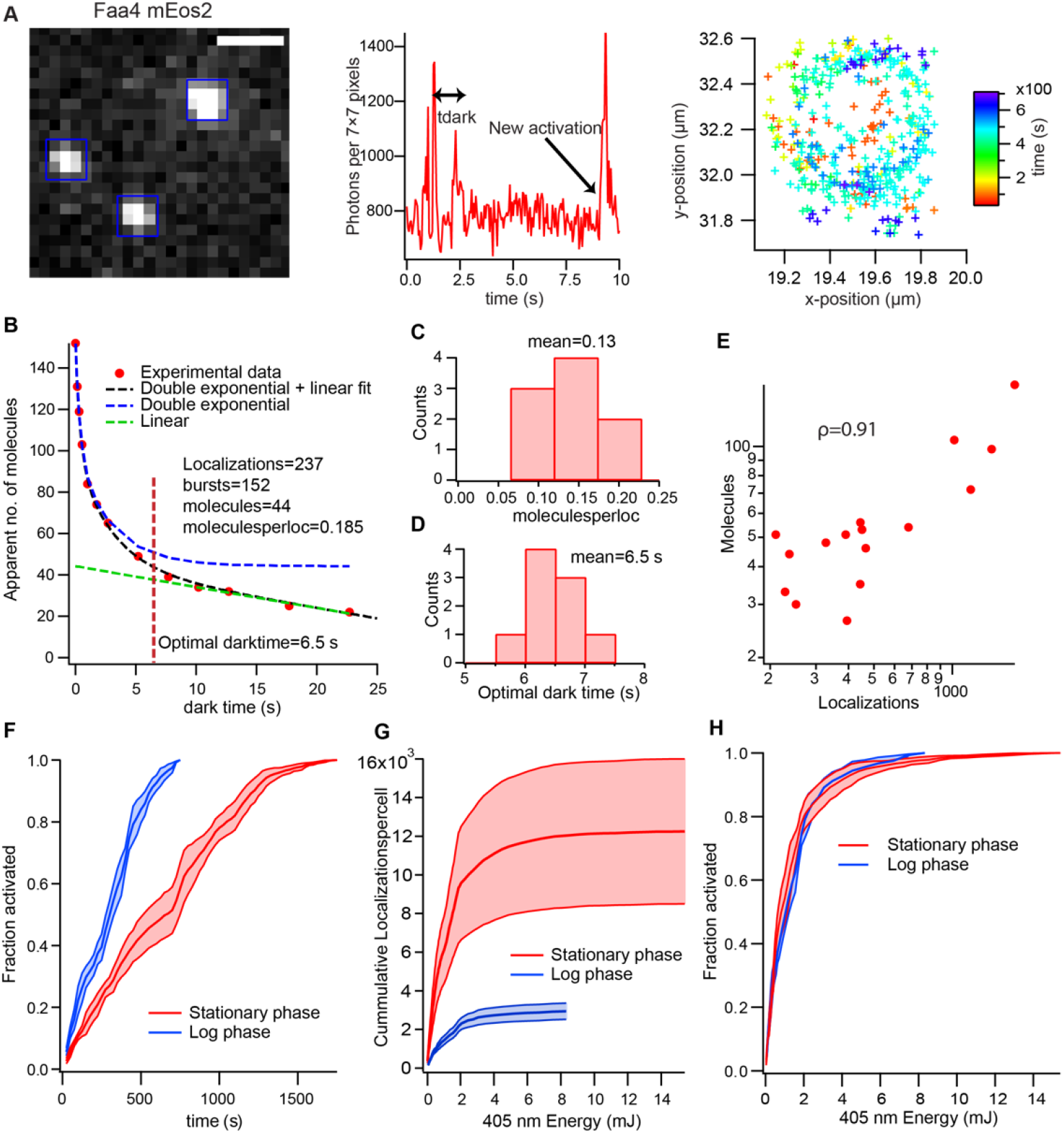
Quantification of Faa4-mEos2 molecule numbers on LDs in living cells. (A) A typical raw data acquisition frame (left) with sparse mEos2 localizations. Intensity trace (middle) of a 7×7 pixels region showing blinking of a mEos2 molecule and a newly excited molecule. mEos2 localizations (right) color-coded by frame number on a LD reveal temporally close fluorescent bursts from single mEos2 molecules dispersed on the surface. The diffusion of single mEos2 molecule during the dark time due to blinking leads to dispersed localizations on the LD surface. (B) Number of apparent blink corrected mEos2 molecules (red dots) as a function of allowed dark time td for a LD. The number of apparent molecule is obtained by grouping localizations on the LD that appear within the specified dark times and within the maximum diameter of LDs of 650 nm. Apparent molecules at td=0s represent the number of fluorescent bursts. Apparent molecules vs dark time show a double exponential decay (dominant at shorter dark times due to blinking) and linear decrease (dominant at longer dark time due to false grouping). The black dotted curve is the fit of the sum of a double exponential decay and a linear function. The blue curve shows only double exponential decay of molecules that converges to actual number of molecules on the LD. The green curve shows linear decrease of molecules due to false linking (shifted in y by the number of molecules to show convergence with the data at long lag times). The optimal dark time (6.5s) balances the over counting fraction from the double exponential part by the undercounting fraction from linear part. The number of molecules per localization (0.185) is obtained from total localizations and the obtained number of blink corrected molecules from the fit at the optimal dark time. (C) Distribution of optimal dark times from N=9 different LDs with varying number of localizations with the mean optimal dark time of 6.5s. (D) Distribution of molecules per localizations for N=9 different LDs with the mean of 0.13±0.04. (E) Number of molecules vs localizations for N=16 LDs shows strong correlation (ρ=0.91). The strong correlation validates the approach of finding the number of detected localizations per mEos2 molecule for blink correction using detected localizations. (F) Fraction of total mEos2 localizations vs data acquisition time under two different activation schemes for cells in the log phase (low Faa4-mEos2 expression level) and stationary phase (high Faa4-mEos2 expression level). Due to the lower expression level of Faa4-mEos2, most molecules in log-phase cells can be imaged faster compared to cells in the stationary phase. (G) The number of localizations per cell (5 cells for log and 4 cells for stationary phase) vs 405 nm energy delivered in the log and stationary phase. Error band represents standard error on mean. (H) Fraction of localized molecules vs 405 nm energy shows that fraction of localized molecules only depends on the delivered 405 nm photo-activation energy. Error band represents standard deviation calculated from N=5 cells for both log and stationary phase. Scale bar: 1μm.

**Suppl.Fig.4:**
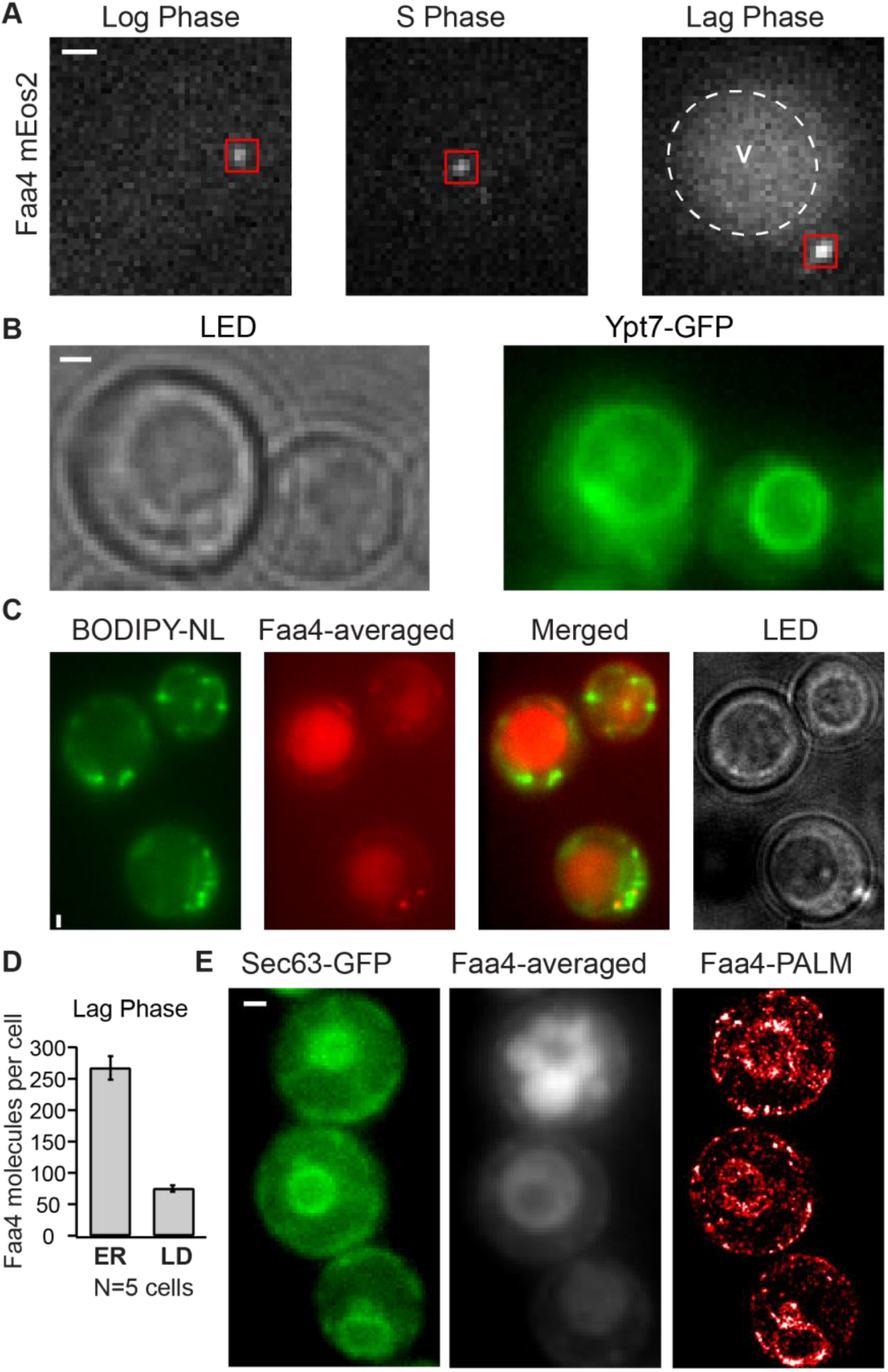
Faa4 is predominately localized to the vacuolar lumen in the lag phase. (A) Single frames of Faa4-mEos2 in the log (left), stationary (middle) and lag (right) phase. Fast diffusion of mEos2 in the vacuole creates a diffuse background fluorescence in the lag phase. (B) Transmitted light image (left) and Ypt7-GFP (right) shows that the darker region in the transmitted light image is the vacuole. (C) BODIPY-NL (left), Faa4-mEos2 avg (middle), merged and LED (right) images show LDs not co-localized with the vacuole. Diffused Faa4 signal comes from dark vacuolar region in the LED image. (D) Quantification of the number of Faa4 molecules in ER and on LDs during the lag phase. (E) The conventional fluorescence of Sec63-GFP (left), Faa4-mEos2 averaged (middle) and Faa4-mEos2 PALM (right) after 30 mins of dilution in fresh medium. A portion of Sec63-GFP signal is seen completely inside Faa4-avg signal from vacuole. Scale Bar: 1 μm

**Supplementary Video1:** Faa4-mEos2 single molecule signal (left) under 561 nm excitation shows blinking mEos2 molecules. Sec63-GFP (right, average of 50 frames) under 488 nm excitation shows ER around the cell periphery and the nucleus. Scale bar: 1 μm.

**Supplementary Video2:** Faa4-mEos2 single molecule signal (left) under 561 nm excitation from cells grown to stationary phase. Nile red signal (right) under 488 nm excitation visualizes a fraction of LDs moving randomly inside vacuole and the other fraction immobile. Faa4-mEos2 signal is also visible from LDs moving randomly inside (top cell). Scale bar: 1 μm.

**Supplementary Video3**: BODIPY-NL signal under 488 nm excitation from a cell grown to stationary phase visualizes a transition of a LD from immobility to random diffusion after it enters the vacuole. Scale bar: 1 μm.

**Supplementary Video4:** Time lapse of averaged Faa4-mEos2 signal (left) and averaged Sec63-GFP signal (right) imaged after adding fresh media for 30 minutes to stationary phase cells. A part of ER (Sec63-GFP) is progressively internalized inside vacuole (Faa4-mEos2) revealing ER-Phagy. Scale bar: 1 μm

## References

Adhikari, S., Moscatelli, J., Smith, E.M., Banerjee, C., and Puchner, E.M. (2019). Single-molecule localization microscopy and tracking with red-shifted states of conventional BODIPY conjugates in living cells. Nat. Commun. 10, 3400.

Annibale, P., Scarselli, M., Kodiyan, A., and Radenovic, A. (2010). Photoactivatable Fluorescent Protein mEos2 Displays Repeated Photoactivation after a Long-Lived Dark State in the Red Photoconverted Form. J. Phys. Chem. Lett. 1, 1506–1510.

Annibale, P., Vanni, S., Scarselli, M., Rothlisberger, U., and Radenovic, A. (2011). Quantitative Photo Activated Localization Microscopy: Unraveling the Effects of Photoblinking. PLoS ONE 6.

Ashrafi, K., Farazi, T.A., and Gordon, J.I. (1998). A role for Saccharomyces cerevisiae fatty acid activation protein 4 in regulating protein N-myristoylation during entry into stationary phase. J. Biol. Chem. 273, 25864–25874.

Bersuker, K., and Olzmann, J.A. (2017). Establishing the lipid droplet proteome: Mechanisms of lipid droplet protein targeting and degradation. Biochim. Biophys. Acta 1862, 1166–1177.

Betzig, E., Patterson, G.H., Sougrat, R., Lindwasser, O.W., Olenych, S., Bonifacino, J.S., Davidson, M.W., Lippincott-Schwartz, J., and Hess, H.F. (2006). Imaging intracellular fluorescent proteins at nanometer resolution. Science 313, 1642–1645.

Black, P.N., and DiRusso, C.C. (2007). Yeast acyl-CoA synthetases at the crossroads of fatty acid metabolism and regulation. Biochim. Biophys. Acta 1771, 286–298.

Brauer, M.J., Huttenhower, C., Airoldi, E.M., Rosenstein, R., Matese, J.C., Gresham, D., Boer, V.M., Troyanskaya, O.G., and Botstein, D. (2008). Coordination of Growth Rate, Cell Cycle, Stress Response, and Metabolic Activity in Yeast. Mol. Biol. Cell 19, 352–367.

De Zitter, E., Thédié, D., Mönkemöller, V., Hugelier, S., Beaudouin, J., Adam, V., Byrdin, M., Van Meervelt, L., Dedecker, P., and Bourgeois, D. (2019). Mechanistic investigation of mEos4b reveals a strategy to reduce track interruptions in sptPALM. Nat. Methods 16, 707–710.

Deschout, H., Shivanandan, A., Annibale, P., Scarselli, M., and Radenovic, A. (2014). Progress in quantitative single-molecule localization microscopy. Histochem. Cell Biol. 142, 5–17.

Durisic, N., Laparra-Cuervo, L., Sandoval-Álvarez, A., Borbely, J.S., and Lakadamyali, M. (2014). Single-molecule evaluation of fluorescent protein photoactivation efficiency using an in vivo nanotemplate. Nat. Methods 11, 156–162.

Faergeman, N.J., Black, P.N., Zhao, X.D., Knudsen, J., and DiRusso, C.C. (2001). The Acyl-CoA synthetases encoded within FAA1 and FAA4 in Saccharomyces cerevisiae function as components of the fatty acid transport system linking import, activation, and intracellular Utilization. J. Biol. Chem. 276, 37051–37059.

Fricke, F., Beaudouin, J., Eils, R., and Heilemann, M. (2015). One, two or three? Probing the stoichiometry of membrane proteins by single-molecule localization microscopy. Sci. Rep. 5, 14072.

Ganesan, S., Sosa Ponce, M.L., Tavassoli, M., Shabits, B.N., Mahadeo, M., Prenner, E.J., Terebiznik, M.R., and Zaremberg, V. (2019). Metabolic control of cytosolic-facing pools of diacylglycerol in budding yeast. Traffic Cph. Den. 20, 226–245.

Greenspan, P., Mayer, E.P., and Fowler, S.D. (1985). Nile red: a selective fluorescent stain for intracellular lipid droplets. J. Cell Biol. 100, 965–973.

Hariri, H., Rogers, S., Ugrankar, R., Liu, Y.L., Feathers, J.R., and Henne, W.M. (2018). Lipid droplet biogenesis is spatially coordinated at ER-vacuole contacts under nutritional stress. EMBO Rep. 19, 57–72.

Hariri, H., Speer, N., Bowerman, J., Rogers, S., Fu, G., Reetz, E., Datta, S., Feathers, J.R., Ugrankar, R., Nicastro, D., et al. (2019). Mdm1 maintains endoplasmic reticulum homeostasis by spatially regulating lipid droplet biogenesis. J. Cell Biol. 218, 1319–1334.

Jacquier, N., Choudhary, V., Mari, M., Toulmay, A., Reggiori, F., and Schneiter, R. (2011). Lipid droplets are functionally connected to the endoplasmic reticulum in Saccharomyces cerevisiae. J. Cell Sci. 124, 2424–2437.

Kiegerl, B., Tavassoli, M., Smart, H., Shabits, B.N., Zaremberg, V., and Athenstaedt, K. (2019). Phosphorylation of the lipid droplet localized glycerol–3–phosphate acyltransferase Gpt2 prevents a futile triacylglycerol cycle in yeast. Biochim. Biophys. Acta Mol. Cell Biol. Lipids 1864, 158509.

Kory, N., Farese, R.V., and Walther, T.C. (2016). Targeting Fat: Mechanisms of Protein Localization to Lipid Droplets. Trends Cell Biol. 26, 535–546.

Kurat, C.F., Natter, K., Petschnigg, J., Wolinski, H., Scheuringer, K., Scholz, H., Zimmermann, R., Leber, R., Zechner, R., and Kohlwein, S.D. (2006). Obese yeast: triglyceride lipolysis is functionally conserved from mammals to yeast. J. Biol. Chem. 281, 491–500.

Lee, S.-H., Shin, J.Y., Lee, A., and Bustamante, C. (2012). Counting single photoactivatable fluorescent molecules by photoactivated localization microscopy (PALM). Proc. Natl. Acad. Sci. U. S. A. 109, 17436–17441.

Listenberger, L.L., Han, X., Lewis, S.E., Cases, S., Farese, R.V., Ory, D.S., and Schaffer, J.E. (2003). Triglyceride accumulation protects against fatty acid-induced lipotoxicity. Proc. Natl. Acad. Sci. U. S. A. 100, 3077–3082.

Loi, M., Raimondi, A., Morone, D., and Molinari, M. (2019). ESCRT-III-driven piecemeal micro-ER-phagy remodels the ER during recovery from ER stress. Nat. Commun. 10.

Manley, S., Gillette, J.M., Patterson, G.H., Shroff, H., Hess, H.F., Betzig, E., and Lippincott-Schwartz, J. (2008). High-density mapping of single-molecule trajectories with photoactivated localization microscopy. Nat. Methods 5, 155–157.

Markgraf, D.F., Klemm, R.W., Junker, M., Hannibal-Bach, H.K., Ejsing, C.S., and Rapoport, T.A. (2014). An ER protein functionally couples neutral lipid metabolism on lipid droplets to membrane lipid synthesis in the ER. Cell Rep. 6, 44–55.

Mejhert, N., Kuruvilla, L., Gabriel, K.R., Elliott, S.D., Guie, M.-A., Wang, H., Lai, Z.W., Lane, E.A., Christiano, R., Danial, N.N., et al. (2020). Partitioning of MLX-Family Transcription Factors to Lipid Droplets Regulates Metabolic Gene Expression. Mol. Cell.

Natter, K., Leitner, P., Faschinger, A., Wolinski, H., McCraith, S., Fields, S., and Kohlwein, S.D. (2005). The spatial organization of lipid synthesis in the yeast Saccharomyces cerevisiae derived from large scale green fluorescent protein tagging and high resolution microscopy. Mol. Cell. Proteomics MCP 4, 662–672.

Olzmann, J.A., and Carvalho, P. (2019). Dynamics and functions of lipid droplets. Nat. Rev. Mol. Cell Biol. 20, 137–155.

Ouahoud, S., Fiet, M.D., Martínez-Montañés, F., Ejsing, C.S., Kuss, O., Roden, M., and Markgraf, D.F. (2018). Lipid droplet consumption is functionally coupled to vacuole homeostasis independent of lipophagy. J. Cell Sci. 131.

Puchkov, E.O. (2010). Brownian motion of polyphosphate complexes in yeast vacuoles: characterization by fluorescence microscopy with image analysis. Yeast Chichester Engl. 27, 309–315.

Puchkov, E.O. (2012). Single yeast cell vacuolar milieu viscosity assessment by fluorescence polarization microscopy with computer image analysis. Yeast Chichester Engl. 29, 185–190.

Puchner, E.M., Walter, J.M., Kasper, R., Huang, B., and Lim, W.A. (2013). Counting molecules in single organelles with superresolution microscopy allows tracking of the endosome maturation trajectory. Proc. Natl. Acad. Sci. U. S. A. 110, 16015–16020.

Romanauska, A., and Köhler, A. (2018). The Inner Nuclear Membrane Is a Metabolically Active Territory that Generates Nuclear Lipid Droplets. Cell 174, 700–715.e18.

Saks, V., Beraud, N., and Wallimann, T. (2008). Metabolic Compartmentation – A System Level Property of Muscle Cells. Int. J. Mol. Sci. 9, 751–767.

Schäfer, J.A., Schessner, J.P., Bircham, P.W., Tsuji, T., Funaya, C., Pajonk, O., Schaeff, K., Ruffini, G., Papagiannidis, D., Knop, M., et al. (2019). ESCRT machinery mediates selective microautophagy of endoplasmic reticulum in yeast. EMBO J. e102586.

Schuldiner, M., and Bohnert, M. (2017). A different kind of love - lipid droplet contact sites. Biochim. Biophys. Acta Mol. Cell Biol. Lipids 1862, 1188–1196.

Seo, A.Y., Lau, P.-W., Feliciano, D., Sengupta, P., Gros, M.A.L., Cinquin, B., Larabell, C.A., and Lippincott-Schwartz, J. (2017). AMPK and vacuole-associated Atg14p orchestrate μ-lipophagy for energy production and long-term survival under glucose starvation. ELife 6.

Swaminathan, R., Hoang, C.P., and Verkman, A.S. (1997). Photobleaching recovery and anisotropy decay of green fluorescent protein GFP-S65T in solution and cells: cytoplasmic viscosity probed by green fluorescent protein translational and rotational diffusion. Biophys. J. 72, 1900–1907.

Tauchi-Sato, K., Ozeki, S., Houjou, T., Taguchi, R., and Fujimoto, T. (2002). The surface of lipid droplets is a phospholipid monolayer with a unique Fatty Acid composition. J. Biol. Chem. 277, 44507–44512.

Thompson, R.E., Larson, D.R., and Webb, W.W. (2002). Precise nanometer localization analysis for individual fluorescent probes. Biophys. J. 82, 2775–2783.

Tsuji, T., Fujimoto, M., Tatematsu, T., Cheng, J., Orii, M., Takatori, S., and Fujimoto, T. (2017). Niemann-Pick type C proteins promote microautophagy by expanding raft-like membrane domains in the yeast vacuole. ELife 6.

Walther, T.C., and Farese, R.V. (2012). Lipid Droplets And Cellular Lipid Metabolism. Annu. Rev. Biochem. 81, 687–714.

Wang, C.-W., Miao, Y.-H., and Chang, Y.-S. (2014a). A sterol-enriched vacuolar microdomain mediates stationary phase lipophagy in budding yeast. J. Cell Biol. 206, 357–366.

Wang, C.-W., Miao, Y.-H., and Chang, Y.-S. (2014b). Control of lipid droplet size in budding yeast requires the collaboration between Fld1 and Ldb16. J. Cell Sci. 127, 1214–1228.

Wang, H., Becuwe, M., Housden, B.E., Chitraju, C., Porras, A.J., Graham, M.M., Liu, X.N., Thiam, A.R., Savage, D.B., Agarwal, A.K., et al. (2016). Seipin is required for converting nascent to mature lipid droplets. ELife 5.

Wu, C.-Y., Roybal, K.T., Puchner, E.M., Onuffer, J., and Lim, W.A. (2015). Remote control of therapeutic T cells through a small molecule-gated chimeric receptor. Science 350, aab4077.

van Zutphen, T., Todde, V., de Boer, R., Kreim, M., Hofbauer, H.F., Wolinski, H., Veenhuis, M., van der Klei, I.J., and Kohlwein, S.D. (2014). Lipid droplet autophagy in the yeast Saccharomyces cerevisiae. Mol. Biol. Cell 25, 290–301.

